# Compound-V formations in shorebird flocks

**DOI:** 10.1101/545301

**Authors:** Aaron J. Corcoran, Tyson L. Hedrick

## Abstract

Animal groups have emergent properties that result from simple interactions among individuals. However, we know little about why animals adopt different interaction rules because of sparse sampling among species. Here, we identify an interaction rule that holds across single and mixed-species flocks of four migratory shorebird species spanning a seven-fold range of body masses. The rule, aligning with a 1-wingspan lateral distance to nearest neighbors in the same horizontal plane, scales linearly with wingspan but is independent of nearest neighbor distance and neighbor species. This rule propagates outward to create a global flock structure that we term the compound-V formation. We propose that this formation represents an intermediary between the cluster flocks of starlings and the simple-V formations of geese and other large migratory birds. Analysis of individual wingbeat frequencies and airspeeds indicates that the compound-V formation may be an adaptation for aerodynamic flocking.

## Introduction

The collective movements of animals—from schooling fish to swarming insects and flocking birds—have long excited intrigue among observers of nature. Collective motion arises as an emergent property of interactions between individuals [reviewed by (Herbert-Read, 2016; Vicsek & Zafeiris, 2012)]. Thus, much attention has been placed on identifying local interaction rules (Ballerini et al., 2008; Herbert-Read et al., 2011; Katz, Tunstrøm, Ioannou, Huepe, & Couzin, 2011; Lukeman, Li, & Edelstein-Keshet, 2010) and how those rules affect group structure and movement (Buhl et al., 2006; Hemelrijk & Hildenbrandt, 2012). However, comparative data across species are still limited, preventing us from testing hypotheses regarding the evolution and diversity of collective movement patterns.

For example, out of hundreds of bird species that fly in groups, most research has focused on starlings (Attanasi et al., 2014; Ballerini et al., 2008; Cavagna et al., 2010), homing pigeons (Nagy et al., 2013; Nagy, Ákos, Biro, & Vicsek, 2010; Pettit, Ákos, Vicsek, & Biro, 2015; Pettit, Perna, Biro, & Sumpter, 2013; Usherwood, Stavrou, Lowe, Roskilly, & Wilson, 2011) and birds that fly in V-formations (Badgerow & Hainsworth, 1981; Cutts & Speakman, 1994; Hummel, 1983; Lissaman & Shollenberger, 1970; Maeng, Park, Jang, & Han, 2013; Portugal et al., 2014; Weimerskirch, Martin, Clerquin, Alexandre, & Jiraskova, 2001). These data indicate that smaller birds fly in relatively dense cluster flocks that facilitate group cohesion and information transfer (Attanasi et al., 2014; Ballerini et al., 2008), whereas larger migratory birds fly in highly-structured V formations (also known as line or echelon formations) that provide aerodynamic and energetic benefits (Lissaman & Shollenberger, 1970; Portugal et al., 2014; Weimerskirch et al., 2001). The species whose flocking behavior have been studied in detail differ in many ways that could be important for flocking including body size, ecology, the frequency of aggregation and its behavioral context. Therefore, it is difficult to conclude based on the available data what factors contribute to birds adopting specific group formations.

We aimed to address this question by collecting three-dimensional (3D) trajectories of the birds in flocks of four shorebird species that have similar ecologies (all forage in large groups in coastal habitats and migrate long distances) but cover a seven-fold range of body mass and two-fold range of wingspan. Our study species include dunlin (*Calidris alpina*, Linnaeus 1758; 56 g, 0.34 m wingspan), short-billed dowitcher (*Limnodromus griseus*, Gmelin 1789; 110 g, 0.52 m wingspan), American avocet (*Recurvirostra Americana*, Gmelin 1789; 312 g, 0.72 m wingspan), and marbled godwit (*Limosa fedoa*, Linnaeus, 1758; 370 g, 0.78 m). Molecular dating indicates that these species diverged from their nearest common ancestor by approximately 50 Mya (Baker, Pereira, & Paton, 2007), providing time for evolutionary diversification of flocking behavior. By comparing group structure of birds across a range of body sizes and comparing our data from that in the literature, we aimed to determine the extent to which flock structure varies across species with different body sizes and ecologies. We employ three approaches: 1) identify local interaction rules by quantifying the relative positions of birds and their nearest neighbors; 2) quantify the degree of spatial structure within flocks; and 3) use measurements of individual speeds and wingbeat frequencies to test whether local or global position within the flock affects flight performance.

Based on existing flock data, we hypothesized that flocks of larger shorebird species would be more structured than smaller species (recapitulating the trend of larger birds flying in highly-structured V formations) and that larger species would also more frequently exhibit aerodynamic formations. Because a previous study showed that flying in a cluster flock is energetically costly in pigeons (Usherwood et al., 2011), we hypothesized that birds flying in the middle and rear of flocks and birds flying closer to their nearest neighbor would have reduced flight performance (lower speed relative to their wingbeat frequency). Surprisingly, we found that all four species studied here fly in a previously-undescribed flock structure that we term the compound-V formation. We propose that this structure is an adaptation for aerodynamic flocking in migratory species, and that ecology is an underappreciated driver of the evolution of avian flocking behavior.

## Results

We reconstructed the 3D trajectories from 18 bird flocks that ranged in size from 189 to 961 individuals and were recorded for 2.4 – 13.2 s at 29.97 frames per second (Figure 1, Table 1). This resulted in 1,598,169 3D position measurements that were used for examining flock structure. Sixteen of the 18 flocks were comprised entirely of a single species. The remaining two flocks were mixed-species flocks of marbled godwits and short-billed dowitchers. Computer vision techniques allowed species identification of individuals in mixed-species flocks based on differences in body size (See Materials and Methods).

**Table 1.** Flock parameters. Values are medians (top) and 10^th^-90^th^ percentiles (bottom) of values extracted at 1-wingbeat intervals from all individuals of each flock. n.n. dist., nearest neighbor distance, values in italics are in wingspan units instead of metric units; n. n. power, exponent of power law fit to distance of 10 nearest neighbors. Wind direction is relative to the overall flight direction where 0° is a pure headwind and 180° a pure tailwind. Note that data are presented separately in consecutive rows for each species in mixed-species flocks (0417-2 and 0417-4).

| Flock | Species | N birds | N Frames | n.n. dist (m) (WS) | n.n. power | Ground speed (m·s<sup>-1</sup>) | Airspeed (m·s<sup>-1</sup>) | Wind speed (m·s<sup>-1</sup>) | Wind direction (deg.) | Z-speed (m·s<sup>-1</sup>) | Turn rate (° s<sup>-1</sup>) |
| --- | --- | --- | --- | --- | --- | --- | --- | --- | --- | --- | --- |
| 0417-2 | Godwit | 286 | 147 | 1.30 1.67<br>0.79, 2.15 | 0.361 | 5.43<br>3.64, 8.66 | 9.36<br>8.04, 10.55 | 5.73 | 5.9 | 1.12<br>0.09, 1.97 | 42.6<br>18.5, 91.8 |
| 0417-2 | Dowitcher | 309 | 147 | 1.17 2.25<br>0.73, 1.89 | 0.385 | 5.30<br>3.56, 8.56 | 9.25<br>7.95, 10.48 | 5.73 | 5.9 | 1.29<br>0.23, 2.08 | 45.8<br>20.1, 97.7 |
| 0417-3 | Godwit | 474 | 278 | 1.81 2.32<br>1.01, 2.94 | 0.428 | 3.33<br>1.72, 6.56 | 7.94<br>6.21, 9.78 | 5.73 | 3.9 | -0.18<br>-1.27, 0.76 | 35.9<br>0.04, 121.2 |
| 0417-4 | Godwit | 802 | 397 | 1.71 2.19<br>1.02, 2.57 | 0.391 | 6.67<br>3.29, 9.57 | 8.84<br>7.69, 10.06 | 5.73 | -312.1 | 0.22<br>-0.78, 1.09 | 16.4<br>5.2, 50.5 |
| 0417-4 | Dowitcher | 74 | 397 | 1.57 2.25<br>0.94, 2.37 | 0.382 | 4.49<br>2.76, 7.61 | 8.69<br>7.63, 9.75 | 5.73 | -312.1 | 0.41<br>-0.60, 1.26 | 22.5<br>7.2, 59.3 |
| 0420-1 | Godwit | 628 | 177 | 1.19 1.53<br>0.60, 1.94 | 0.409 | 7.05<br>5.14, 9.85 | 10.62<br>8.57, 13.42 | 4.06 | 5.5 | -0.26<br>-1.83, 1.02 | 24.5<br>8.1, 95.4 |
| 0420-2 | Godwit | 304 | 147 | 1.54 1.97<br>0.84, 2.37 | 0.431 | 8.92<br>7.08, 11.55 | 10.55<br>9.65, 11.84 | 4.06 | 54.8 | -0.07<br>-0.95, 0.67 | 13.1<br>4.3, 46.7 |
| 0427-2 | Dowitcher | 354 | 397 | 1.08 2.08<br>0.62, 1.99 | 0.424 | 6.22<br>3.07, 9.76 | 9.17<br>5.50, 12.80 | 3.50 | 22.8 | 0.56<br>-0.90, 1.79 | 22.6<br>7.5, 71.0 |
| 0427-3 | Dowitcher | 391 | 218 | 1.14 2.19<br>0.61, 2.05 | 0.463 | 10.87<br>8.64, 13.30 | 11.07<br>8.91, 14.44 | 3.50 | 71.5 | 0.33<br>-2.10, 2.15 | 30.3<br>9.7, 91.0 |
| 0427-5 | Dowitcher | 510 | 171 | 1.23 2.37<br>0.70, 1.98 | 0.421 | 5.31<br>4.23, 6.49 | 9.52<br>8.56, 10.49 | 4.60 | 14.1 | 0.73<br>-0.24, 1.52 | 24.9<br>8.8, 75.3 |
| 1230-1 | Dunlin | 351 | 198 | 1.08 3.17<br>0.58, 1.78 | 0.465 | 6.99<br>6.04, 7.85 | 6.66<br>5.95, 7.54 | 1.09 | 102.7 | -0.23<br>-0.52, 0.09 | 29.1<br>10.5, 50.0 |
| 1230-2 | Dunlin | 578 | 75 | 0.80 2.35<br>0.47, 1.29 | 0.387 | 6.61<br>6.00, 7.30 | 7.39<br>6.80, 8.01 | 1.09 | 41.6 | -0.13<br>-0.57, 0.32 | 15.6<br>4.9, 40.5 |
| 1230-3 | Dunlin | 477 | 125 | 0.86 2.53<br>0.49, 1.37 | 0.392 | 6.71<br>5.85, 7.74 | 7.55<br>6.78, 8.51 | 1.09 | 34.3 | 0.11<br>-0.50, 0.84 | 26.4<br>7.6, 69.3 |
| 1230-4 | Dunlin | 189 | 73 | 0.89 2.62<br>0.50, 1.50 | 0.503 | 8.28<br>7.46, 9.70 | 7.49<br>6.69, 9.17 | 1.09 | 132.5 | -0.03<br>-0.57, 0.45 | 30.2<br>7.8, 82.2 |
| 0101-1 | Dunlin | 289 | 350 | 1.59 4.68<br>0.69, 3.04 | 0.490 | 7.55<br>5.96, 8.82 | 7.30<br>6.21, 8.84 | 1.63 | 89.9 | -0.70<br>-1.28, -0.08 | 18.8<br>5.6, 41.6 |
| 0101-3 | Dunlin | 961 | 188 | 0.92 2.71<br>0.52, 1.49 | 0.416 | 8.61<br>7.70, 9.42 | 7.75<br>7.11, 8.50 | 1.63 | 117.5 | 0.02<br>-0.33, 0.36 | 15.6<br>4.3, 36.8 |
| 0101-4 | Dunlin | 269 | 340 | 1.00 2.94<br>0.35, 2.11 | 0.451 | 6.56<br>4.96, 8.13 | 7.73<br>6.28, 8.89 | 1.63 | 38.6 | -0.14<br>-0.70, 0.30 | 27.5<br>6.7, 76.1 |
| 1220-1 | Avocet | 321 | 90 | 1.09 1.51<br>0.70, 1.68 | 0.429 | 6.03<br>4.55, 7.57 | 8.21<br>6.74, 9.53 | 2.39 | 8.5 | 0.33<br>-0.98, 0.90 | 27.0<br>10.3, 51.2 |
| 1220-2 | Avocet | 321 | 245 | 1.19 1.65<br>0.72, 1.89 | 0.432 | 6.96<br>5.26, 9.22 | 8.04<br>7.04, 9.12 | 2.39 | 1.7 | 0.10<br>-1.49, 0.74 | 31.2<br>11.7, 64.6 |
| 1220-3 | Avocet | 281 | 280 | 1.30 1.81<br>0.79, 2.10 | 0.472 | 7.50<br>5.32, 8.93 | 7.96<br>6.87, 8.91 | 2.39 | 33.1 | 0.33<br>-0.62, 0.83 | 25.7<br>6.6, 58.3 |

**Figure 1.**
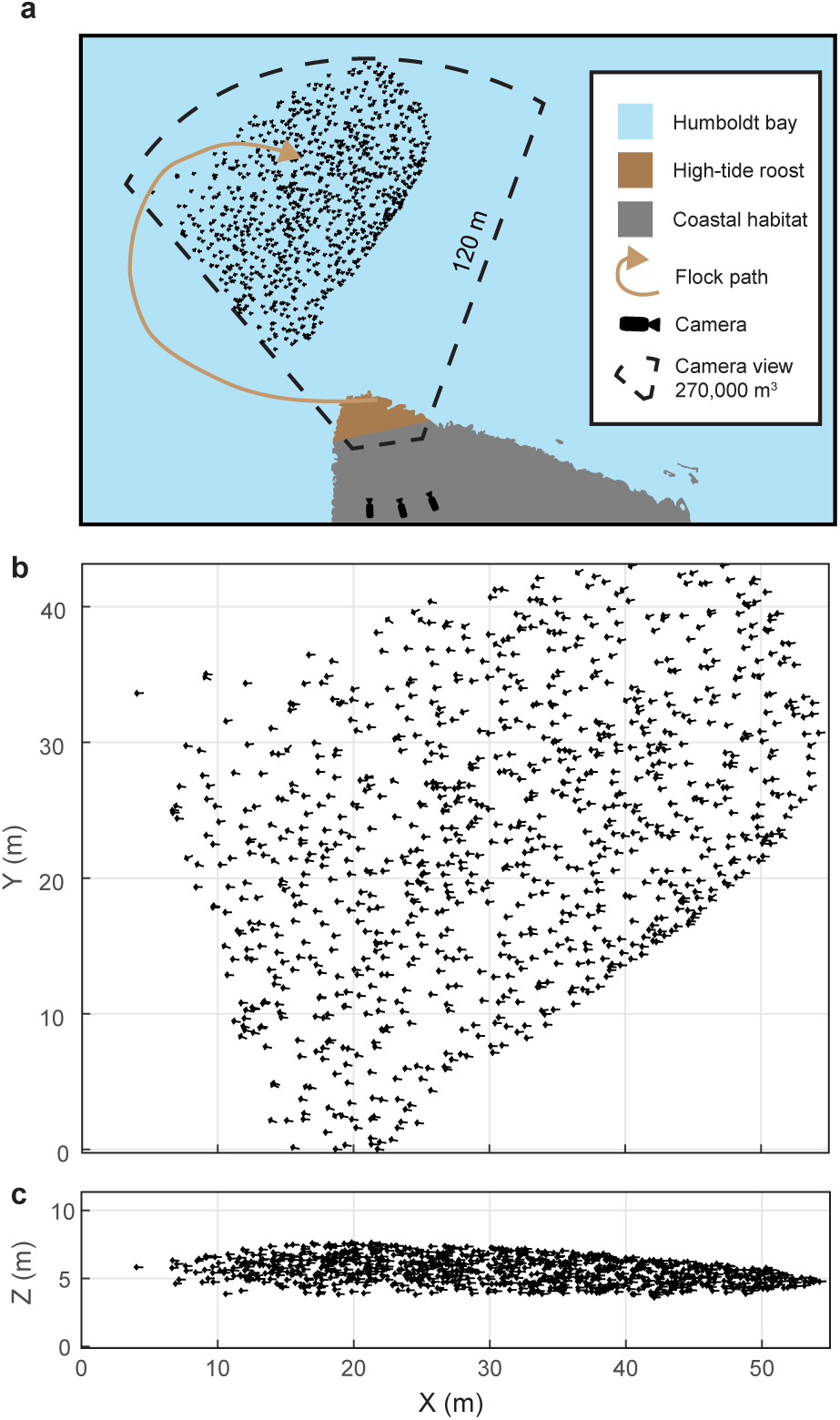
Shorebird flock recording. (a) Multi-camera videography was used to reconstruct 3D trajectories of shorebirds flying near high-tide roosts in Humboldt Bay, California. (b) Overhead and (c) profile views of an example flock. Symbol sizes reflect actual scales for birds with outstretched wings.

### Nearest neighbor alignment

We examined flock structure by quantifying the position of each bird with respect to its nearest neighbor. We used modal values to characterize typical neighbor positions because position distributions were skewed as a result of values being cropped at zero. In all flocks, nearest neighbors flying within the same horizontal plane (an elevation slice of ± 1 wingspan, mean of 56% of nearest neighbors across all flocks; range 35-76%) exhibited a distinctly peaked distribution where modal neighbor position was offset both in front back and lateral distance (Figure 2a, b). In contrast, nearest neighbor birds flying outside the horizontal elevation slice of ± 1 wingspan were distributed randomly with a peak directly above or below the focal bird (Figure 2c, d). This indicates that shorebirds adopt alignment rules for neighbors flying within their same elevation slice.

**Figure 2.**
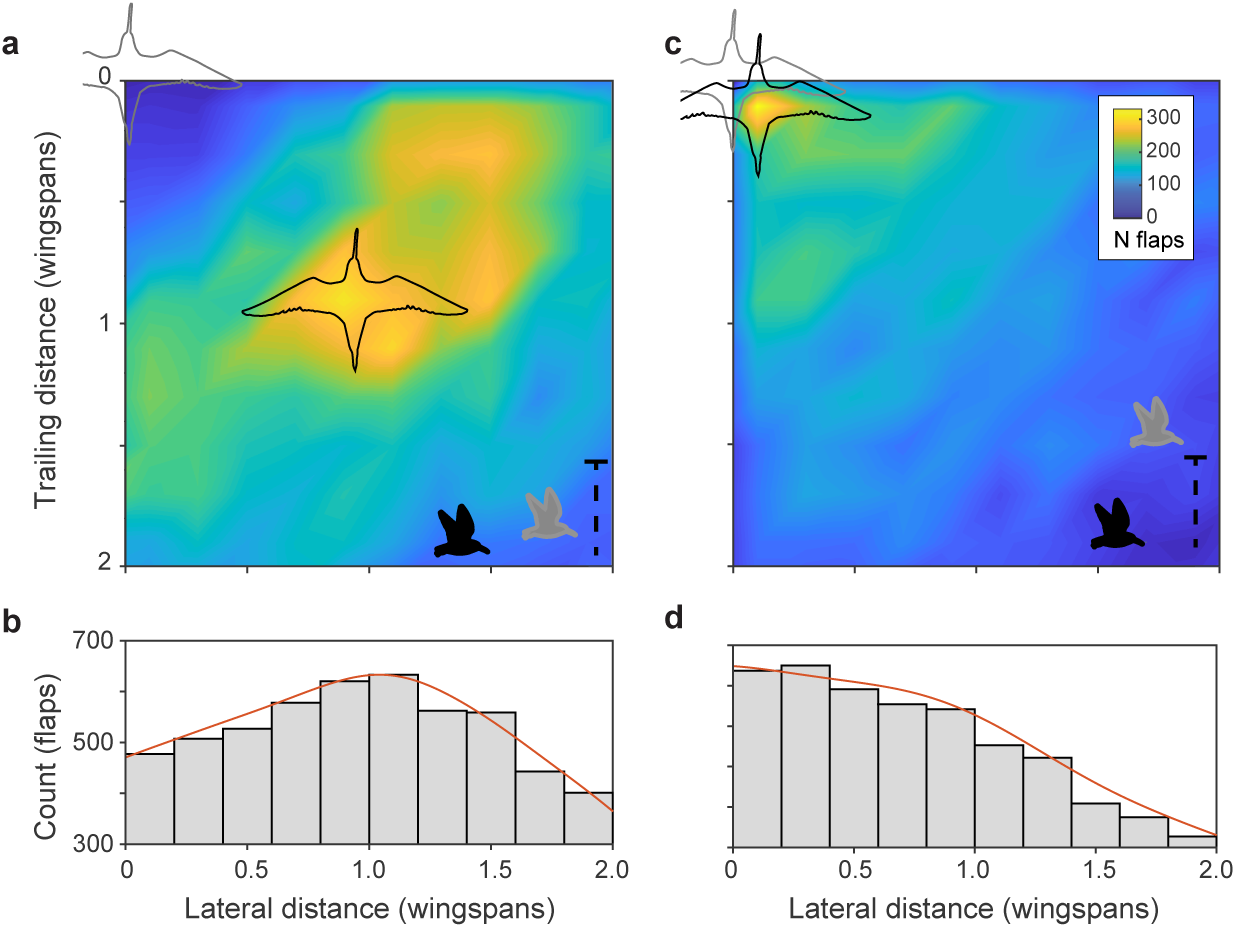
Within-flock positioning. (a, b) Histograms of nearest-neighbor alignment for birds flying within ± 1 wingspan of elevation (godwit flock 0420-1) show a distinctive peak at a trailing distance and lateral distance of approximately 1 wingspan; focal birds are shown in light gray and nearest neighbors in black. Inset bird silhouettes show profile views of the birds’ relative flight elevations. (c, d) Histograms of nearest-neighbor alignment for birds flying outside ± 1 wingspan of elevation for the same flock show a largely random distribution with a modal location of nearly straight above or below the focal bird.

Both nearest-neighbor lateral distance and front-back distance differed among flocks and species (Figure 3a). Species wingspan strongly predicted modal lateral neighbor position (linear regression, slope = 0.85, R^2^ = 0.93, F = 228.29, P < 0.0001). Wingspan also predicted front-back distance (slope = 0.70, R^2^ = 0.86, F = 99.57, p < 0.0001), although less strongly than lateral distance. After scaling alignment positions to wingspan (i.e., dividing neighbor distances by species wingspan), a distinctive pattern emerges (Figure 3b). Specifically, the flocks adopted a modal lateral distance of approximately 1 wingspan (mean 1.04, range 0.88 – 1.24 wingspans). This non-dimensionalized lateral distance had a weak inverse relationship to species wingspan (linear regression, slope = −0.37, R^2^ = 0.37, F = 9.38, P = 0.007) and was not related to flock density (i.e. nearest neighbor distance, non-dimensionalized by wingspan; linear regression, R^2^ = 0.07, F = 1.15; P = 0.30). Non-dimensionalized trailing distance was inversely proportional to species wingspan (linear regression, slope = - 0.40, R^2^ = 0.33, F = 7.93, P = 0.012) and increased with non-dimensional flock density (linear regression, slope = 0.13, R^2^ = 0.58, F = 22.39; P = 0.0002). In summary, across all four species, shorebirds adhere to a non-dimensional spacing rule of aligning to neighbors with a lateral offset of approximately one wingspan while allowing trailing distance to vary with flock density.

**Figure 3.**
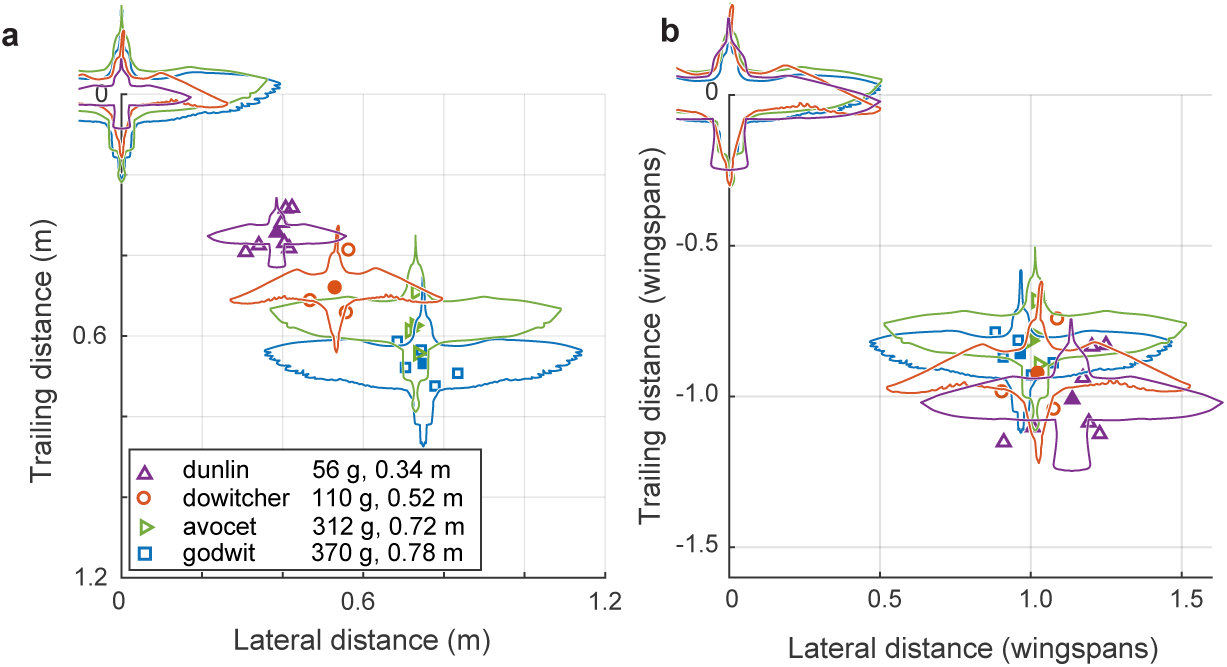
Modal positioning among flocks and species. (a) Summary of modal neighbor position for nearest neighbors within a ± 1 wingspan in single-species flocks of all four species, depicted in absolute metric distance and (b) the same data plotted in distances relative to the wingspan of each species. Open symbols indicate modal neighbor positions for individual flocks. Closed symbols and silhouettes show the average position for each species.

Data from mixed-species flocks of godwits and dowitchers further support the non-dimensional nature of the lateral spacing rule within individual flocks. Both dowitchers and godwits adjusted their lateral spacing depending on the species of their neighbor (Figure 4). Godwits following conspecifics had a modal lateral spacing of 0.76 m, or 0.97 godwit wingspans. When following the smaller dowitchers, godwits reduced the modal lateral distance to 0.60 m or 0.92 wingspans when calculated using the average wingspan of dowitchers and godwits (Mann-Whitney *U* = 525684; *n*_1_ = 1034; *n*_2_ = 81; P = 0.0004). Dowitchers following conspecifics flew with a modal lateral distance of 0.51 m, or 0.98 dowitcher wingspans. When following the larger godwits, dowitchers increased the modal lateral distance to 0.58 m or 0.89 average wingspans (Mann-Whitney U test; U = 66341; *n*_1_ = 743; *n*_2_ = 149; P < 0.0003).

**Figure 4.**
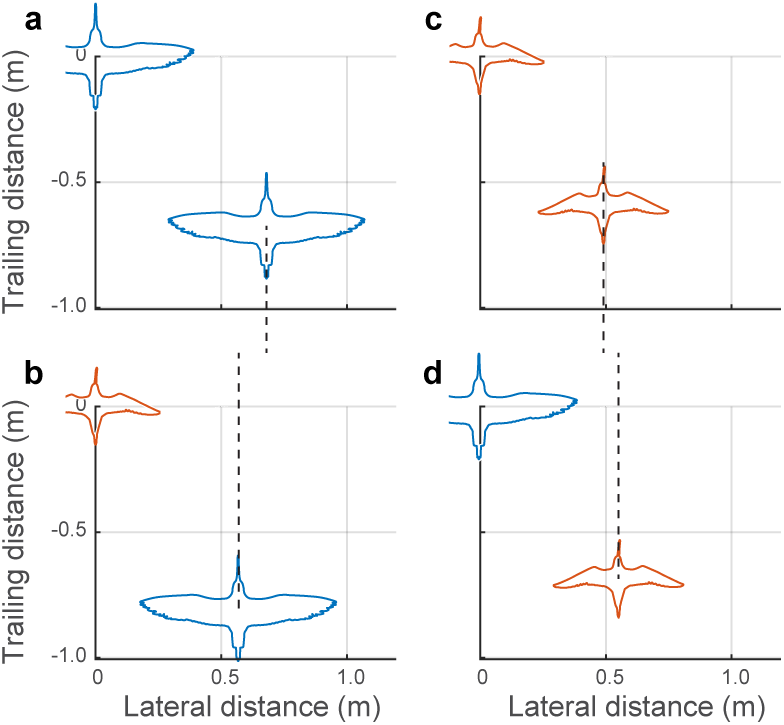
Positioning in mixed-species flocks. Data from mixed species flocks show that birds adjust their lateral spacing depending on the species (and size) of their nearest leading neighbor. (a) Godwits following conspecifics adopt a larger lateral distance than (b) godwits following the smaller dowitchers. (c) Dowitchers following conspecifics use a shorter lateral distance than (d) dowitchers following the larger godwits (See Results for statistics). These results support the hypothesis that shorebirds adopt a lateral spacing rule that is dependent on the size of their leading neighbor. Dashed lines are provided to facilitate comparison of modal lateral positions between (a) and (b) and between (c) and (d).

### Comparison of simple- and compound-V formations

While recording the larger cluster flocks, we also recorded four godwit simple-V formations having between 16-44 individuals and that were recorded for between 42-211 frames (Figure 5). Here we compare the positioning of godwits in simple and compound-V formations. In both cases, nearest neighbors were most commonly in the same horizontal plane (mean of 61% in godwit cluster flocks, 97.9% in godwit simple-V formations), defined as extending 1 wingspan above and below the focal bird, and that the follower is positioned over a narrow lateral range and wider range of trailing distances (Figures 3b, 5b). The modal lateral position in the simple-V formations was slightly less (mean of 0.8 wingspans) than in the compound-V formations where the mean modal lateral position among godwit flocks was 0.96 wingspans (Generalized Linear Model with terms for flock and simple *vs* compound-V formation; P < 0.0001). The modal trailing distance in simple-V formation was 0.50 wingspans; in compound-V formations of godwits, the mean of modal trailing distances was 0.86 wingspans.

**Figure 5.**
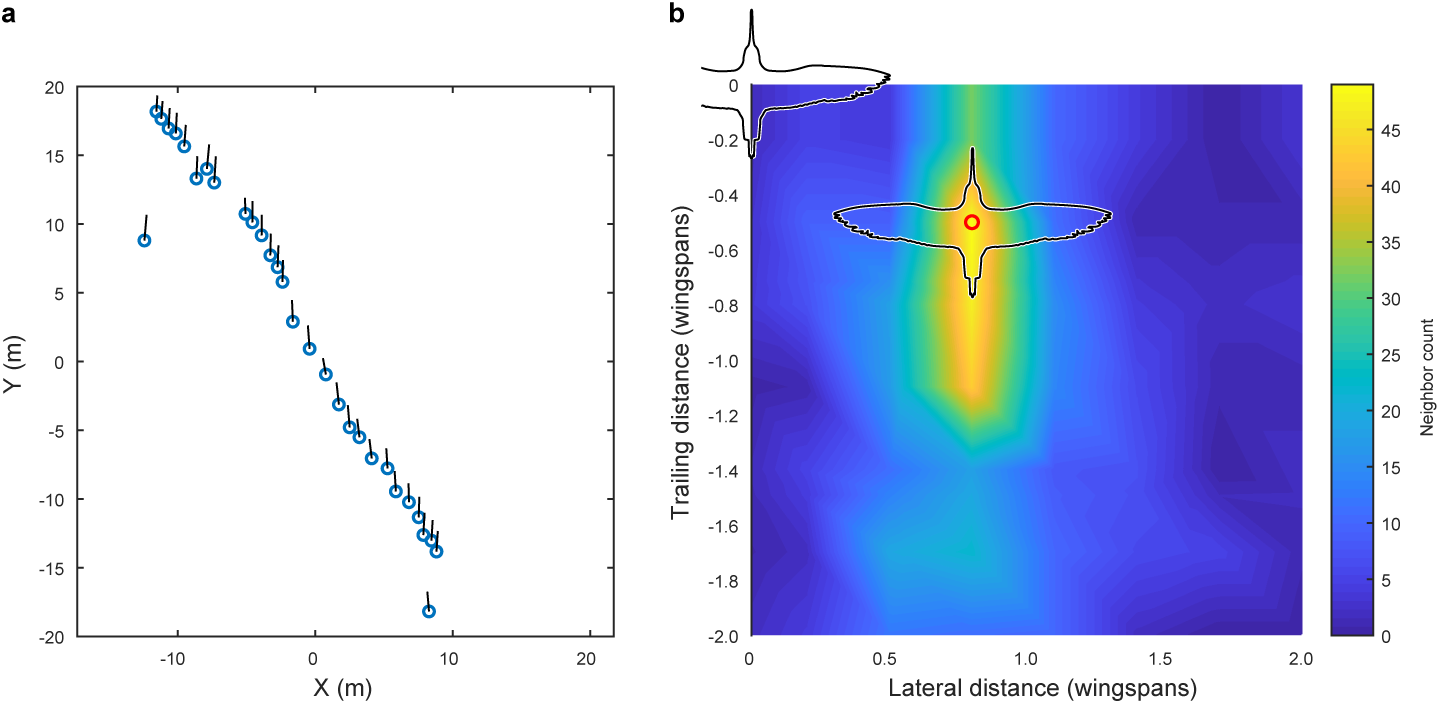
Godwit simple-V formation. Incidental to our cluster flock recordings, we also recorded several instances of godwits flying in a simple-V, echelon or line formation, the largest of these examples is shown here. Panel (a) provides an overhead view of the flock; average flight direction is along the positive Y axis; blue circles show bird positions and black lines are 2D velocity vectors. All birds are within a ± 1 wingspan horizontal slice. Panel (b) shows the relative location of nearest neighbors; the modal location (red circle) was at a displacement of 0.8 wingspans lateral and 0.5 wingspans trailing distance. As the histogram in (b) shows, trailing position was more varied than lateral position. Wind speed was low (< 2 m s^−1^) according to weather station data and wind speed estimated from the ground speed and flight direction of the birds.

### Extended flock structure

We next examined how individual neighbor alignment rules relate to flock structure. We measured the angular distribution of neighbors at distances of 2, 4, 6, and 8 wingspans and at the maximum distance where half of the flock remains in the flock’s core (Range 5.8-24.3 wingspans). This last measure was used as a proxy for whole flock structure while avoiding edge effects (See Methods). At a distance of 2 wingspans, flocks were consistently asymmetrical with trailing birds more frequently flying to the left of their leading neighbors in 12 of 18 flocks and to the right of their nearest leading neighbors in the remaining six flocks; this asymmetry persisted at all distances within the flock (Figure 6a), including the overall flock shape (Figure 6b). The direction of asymmetry was independent of relative camera viewing direction and flock turning direction but was positively correlated to relative wind direction (Table 2).

**Table 2.** Flock orientation. Tests of the relationship between flock left-right orientation (Figure 6) and environmental factors

| Variable | Test | N | R<sup>2</sup> | t/F | P |
| --- | --- | --- | --- | --- | --- |
| Wind direction | Circular correlation | 18 | 0.29 | 2.29 | 0.02 |
| Turn direction | Linear regression | 18 | 0.00 | 0.04 | 0.83 |
| Camera direction | Circular correlation | 18 | 0.02 | 0.61 | 0.54 |

**Figure 6.**
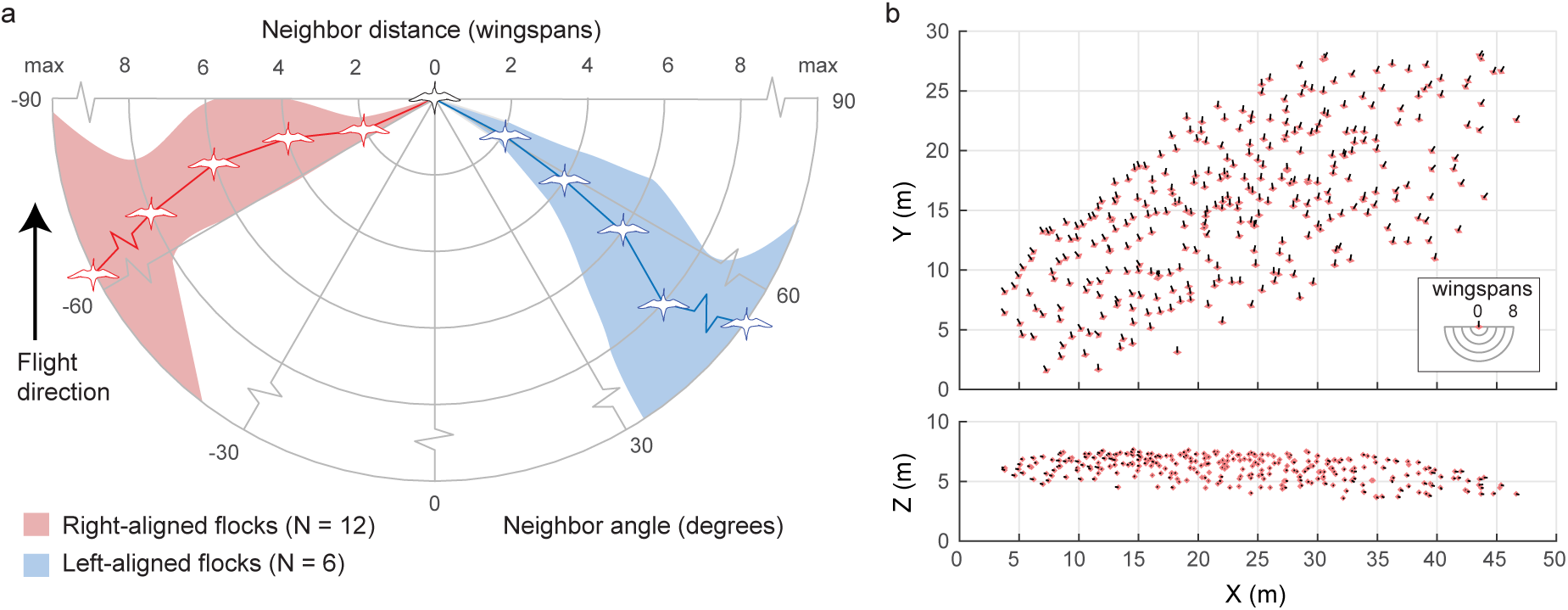
Extended flock structure. (a) Polar plot showing mean neighbor angle for right-aligned and left-aligned flocks at a range of distances. Shaded regions show 95% confidence intervals. (b) Overhead and profile view of an example right-aligned flock (avocet flock 1220-2). Note the many echelon formations aligned from back left to front right and the overall shape of the flock. The inset shows scale in wingspans.

### Flock biomechanics

We quantified several biomechanically relevant parameters from individual birds in flocks, including ground speed, air speed, ascent or descent speed, wingbeat frequency and flapping phase. We created statistical models to predict wingbeat frequency and airspeed from local and global flock position and other flight parameters. While speeds were measured for all individuals, flapping frequency and phase were only available from 6 flocks where birds were sufficiently close to cameras for wingbeat measurements. We examine only data where wingbeat and airspeed data were available (N = 3,306 individuals). We were also unable to measure flapping parameters from Dunlin, the smallest species recorded here.

We observed several individual and flock effects on flight speed and wingbeat frequency (Table 3). As expected, different species flew with different characteristic flapping frequencies and speeds, and climbing flight was associated with an increase in flapping frequency. Birds flying near the front of the flock along the direction of travel (birds were given a continuous index with 0 being the frontmost and 1 the rearward-most position) flew faster and with a lower flapping frequency than those near the rear. Birds flying near the edge of the flock also flew faster than those in the middle. Higher flapping frequencies were correlated with slower flight, potentially reflecting the influence of a range of body sizes with larger individuals flapping more slowly while also flying faster (Pennycuick, 1990). Because our data are among individuals rather than among speeds (or frequencies) for an individual, results are not expected to mirror classic U-shaped flight power curve predictions (Pennycuick, 1968). Birds flying within the predicted range of locations for aerodynamic interaction (0.7-1.5 wingspans lateral distance and within 2 wingspans overall distance of leading neighbor, coded as “Aerodynamic neighbor” in Table 3) flew faster than expected after controlling for the other effects described above (Figure 7). However, positioning in this aerodynamic interaction region had no effect on flapping frequency (Table 3).

**Table 3.** Flock biomechanics. n.n., nearest neighbor; only defined when n.n. is leading the focal bird. Godwit and avocet are dummy variables coding species differences relative to dowitchers. Nearest neighbor species is coded −1 for a smaller neighbor, 0 for same species, 1 for larger neighbor. Flock position is scaled from 0 (front) to 1 (back). Aerodynamic neighbor was coded 1 for birds flying with 0.7-1.5 wingspans lateral distance and within 2 wingspans distance from their nearest leading neighbor, 0 otherwise. Models were selected using Bayesian information criteria. Other predictors that were considered, but not included in the final models were ground speed and apparent airspeed.

| Wingbeat frequency predictors | Estimate | s.e. | t | d.f. | P |
| --- | --- | --- | --- | --- | --- |
| Intercept (dowitchers) | 8.82 | 0.132 | 66.5 | 2817 | < 0.00001 |
| Godwit | -2.19 | 0.035 | -63.5 | 2817 | < 0.00001 |
| Avocet | -1.91 | 0.040 | -47.5 | 2817 | < 0.00001 |
| Airspeed (m s<sup>-1</sup>) | -0.05 | 0.013 | -4.3 | 2817 | 0.00002 |
| Flock position | 0.13 | 0.032 | 4.0 | 2817 | 0.00006 |
| n.n. distance (wingspans) | -0.03 | 0.010 | -3.2 | 2817 | 0.00153 |
| n.n. species | -0.45 | 0.038 | 11.9 | 2817 | < 0.00001 |
| Z-speed (m s<sup>-1</sup>) | 0.29 | 0.019 | 15.1 | 2817 | < 0.00001 |
| Airspeed predictors |  |  |  |  |  |
| Intercept (dowitchers) | 10.69 | 0.225 | 47.5 | 2832 | < 0.00001 |
| Godwit | -0.25 | 0.067 | -3.7 | 2832 | 0.00022 |
| Avocet | -1.51 | 0.067 | -22.6 | 2832 | < 0.00001 |
| n.n. distance (wingspans) | -0.08 | 0.015 | -5.2 | 2832 | < 0.00001 |
| Edge distance (wingspans) | -0.03 | 0.003 | -10.3 | 2832 | < 0.00001 |
| Wingbeat frequency (Hz) | -0.12 | 0.025 | -5.0 | 2832 | < 0.00001 |
| Flock position | -0.24 | 0.045 | -5.4 | 2832 | < 0.00001 |
| Aerodynamic neighbor | 0.23 | 0.040 | 5.7 | 2832 | < 0.00001 |

**Figure 7.**
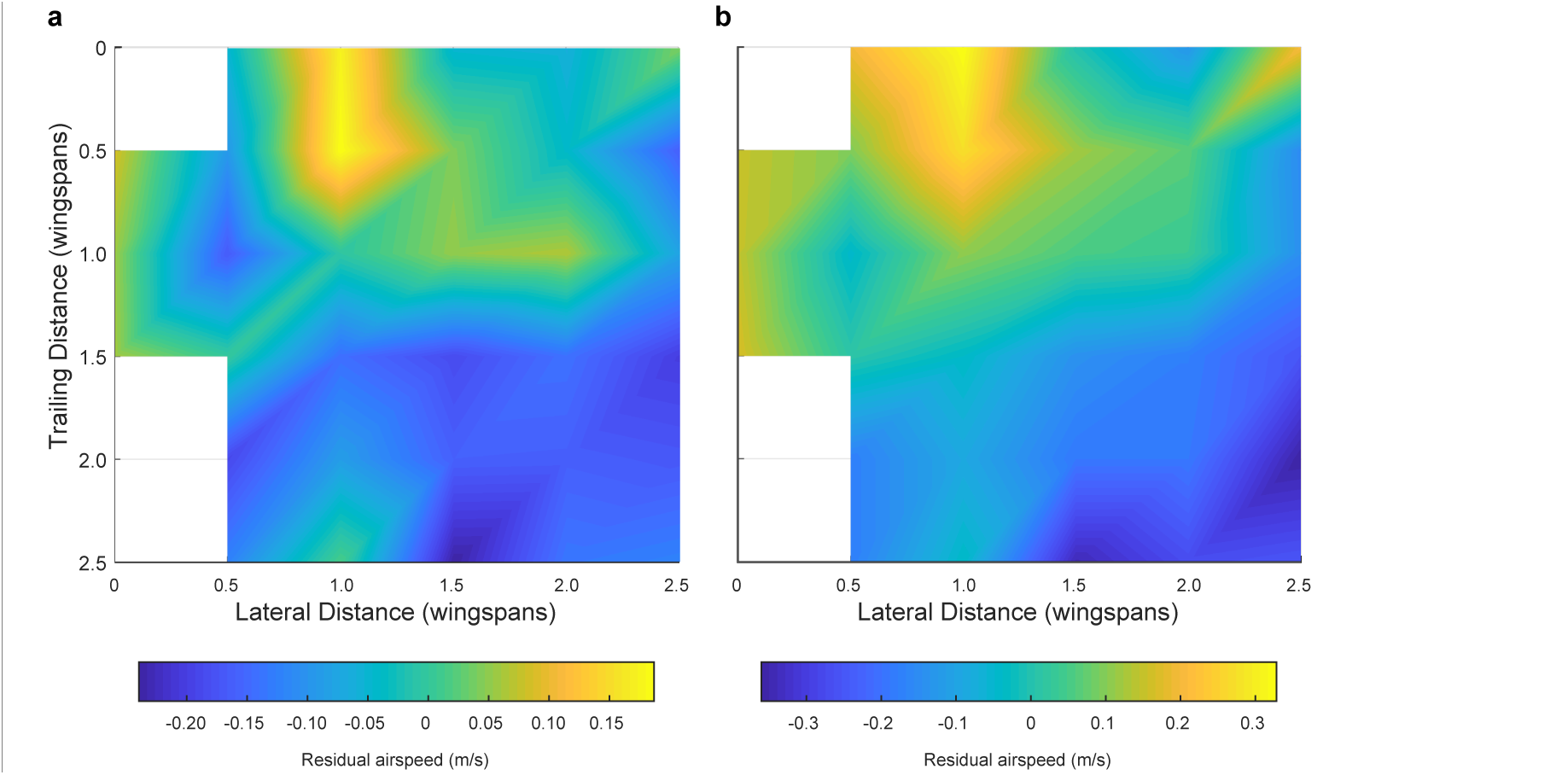
Effect of positioning for aerodynamic interaction. Here we show the effect of neighbor position on flight speed. Panel (a) shows the flight speed residuals after accounting for species, flapping frequency, distance from flock edge, nearest neighbor distance in terms of wingspans and overall position along the length of the flock. Panel (b) shows the flight speed residuals after accounting only for species and flapping frequency. White spaces in the heat map are bins with fewer than 20 samples, out of 2848 possible in (a) and 3306 possible in (b). Both analyses reveal a broadly similar pattern, where the positive effect of neighbor position on flight speed is strongest at a 1 wingspan lateral displacement and a trailing distance of 0 to 0.5 wingspans. This pattern cannot be generated by trailing birds passing leaders because the roles reverse after passing occurs, leaving no net speed difference.

We examined the cluster flock data for evidence of flapping synchronization by examining the temporal and spatial phase offset between pairs of nearest neighbors where synchronous wingbeat frequency data were available for at least 20 frames (See materials and methods). We found no evidence for temporal (Rayleigh test; N = 117; Z = 1.98; P = 0.14) or spatial wingbeat synchronization (Rayleigh test; N = 117; Z = 1.28; P = 0.27) in the compound-V formation shorebird flocks. We performed the same tests on the simple-V formation of godwits and also found no support for phasing relationships (Rayleigh test; temporal phasing N = 39; Z = 0.09; P = 0.90; spatial phasing; N = 39; Z = 0.46, P = 0.63).

## Discussion

Here, we report on the first cross-species analysis of bird flocking behavior. Based on previous studies, we predicted that larger species flocks of four shorebird species covering a 7-fold range in body mass. The simple alignment rule produces a flock structure that can be observed at all spatial scales within the flock, including overall flock shape (Figure 6). This is in contrast to other flocks, such as starlings, where structure is only observed within each neighbor’s six nearest neighbors, equal to 1.2-2.7 wingspans (Ballerini et al., 2008). Our data do show the shorebird global flock alignment is responsive to local wind conditions (Table 2), and future work exploring this interaction may allow identification of the mechanism governing the overall alignment.

We propose that the alignment rule observed in compound-V formations is an adaptation for individuals to gain aerodynamic benefits from flying in the upwash generated by wingtip vortices of their leading neighbors. Both theoretical (Badgerow & Hainsworth, 1981; Hummel, 1983; Lissaman & Shollenberger, 1970; Maeng et al., 2013) and empirical research (Portugal et al., 2014; Weimerskirch et al., 2001) has provided support for the hypothesis that birds flying in simple-V formations gain aerodynamic and energetic benefits. Birds in compound-V formations adopt a similar alignment rule to that observed in simple-V formations. Specifically, in both cases birds fly with a lateral offset of approximately one wingspan while allowing trailing distance to vary (Figures 3 and 5). Compared to simple-V formations, compound-V formations allow higher flock densities, which should allow more rapid information transfer (Attanasi et al., 2014), larger flock sizes, and improved predator defense (Powell, 1974). The compound-V alignment rule contrasts with that observed in starlings (Ballerini et al., 2008), homing pigeons (Pettit et al., 2013) and chimney swifts (Evangelista, Ray, Raja, & Hedrick, 2017), whose alignment rules prevent alignment for beneficial aerodynamic interactions.

Because birds in simple and compound-V formations adopt similar neighbor alignment rules, alternative hypotheses for simple-V formations might apply to compound-V formations. These include collision avoidance and information transfer (Dill, Holling, & Palmer, 1997). Collision avoidance is a plausible hypothesis for simple-V formations because they theoretically permit birds to keep all neighbors out of their direct path of travel. This is not the case for compound-V formations, where many birds are flying in front of and behind one another (Figure 1b; Figure 6b). The problem of collision avoidance is exacerbated in compound-V formation because birds tend to fly in the same horizontal plane. A better strategy for collision avoidance is to fly in a three-dimensional shape, such as that observed in starlings (Ballerini et al., 2008) and swifts (Evangelista et al., 2017). Finally, even in the simple-V formation recorded here (Figure 5) birds flew with approximately 20% of wingspan overlap and so did not have an entirely clear forward path. Thus, collision avoidance appears to be an unlikely explanation for the structuring of both compound-V and simple-V formations.

Simple and compound-V formations might also be structured to maximize the observability of neighbors, facilitating information transfer by helping birds detect and respond to changes in neighbor speed or direction and improving flock cohesiveness by allowing information to propagate through the flock more quickly. Dill and colleagues (Dill et al., 1997) proposed that birds in V formation should maximize measurement of neighbor movements by aligning at a 35.3 degree angle (relative to the direction of travel), or alternatively maximize measurement of neighbor speed by aligning at a 63.4 degree angle. The shorebird flocks examined here had modal neighbor position alignment angles ranging from 33.7 to 51.8 with an average of 41.2 degrees. Neither this mean angle, nor the nearly 20-degree range in alignment angle is consistent with Dill’s hypotheses or others calling for a single optimal alignment angle, and our finding (see above) showing that lateral spacing is uncorrelated with flock density whereas trailing spacing increases with decreasing density shows that the shorebird flocks are more organized in lateral distance than in trailing distance or alignment angle. Thus, hypotheses calling for organization based on alignment angle, whether to maximize information transfer or to keep lead birds in the visual fovea of trailing neighbors in a V formation (Badgerow & Hainsworth, 1981) are not well supported by our results.

Analysis of airspeeds and wingbeat frequencies of flocking shorebirds provides further support for the aerodynamic alignment hypothesis. Birds flying in positions where beneficial aerodynamic interactions are predicted to occur flew faster than expected after controlling for other factors (aerodynamic neighbor term in Table 3). This should produce a reduced cost of transport, assuming there are no unmeasured compensating factors such as a simultaneous increase in stroke amplitude. This result could explain why larger groups of shorebirds fly faster than smaller groups (Hedenström & Åkesson, 2017), as the proportion of birds gaining a speed advantage should increase with flock size.

Theoretically, if birds seek to minimize their cost of transport they should slow down when experiencing a reduction in induced power costs (Hummel, 1983). This is because induced power costs decrease with increasing speed while costs due to drag increase with increasing speed. Thus, a reduction in induced power costs is best taken advantage of by slowing down to reduce drag costs. However, slowing down might disrupt the formation, since the lead bird experiences no benefit and might maintain normal speed. To remain in formation, trailing birds might reduce their power output but maintain speed by reducing flapping frequency or amplitude. A reduction in wingbeat frequency was seen in the simple-V formation flight of pelicans (Weimerskirch et al., 2001), providing some support for this strategy although no similar trend was reported for ibis (Portugal et al., 2014). We also find no reduction in flapping frequency for birds positioned where aerodynamic interactions are expected to be strongest (Table 3). Thus, if shorebirds are gaining an aerodynamic benefit from flying in a neighbor’s upwash, they are not reducing their power output, but instead fly at greater speed.

Speed differences within a flock could be a problem for maintaining flock structure. However, outside the nearest-neighbor alignment relationship, the shorebird flocks appear loosely structured with space between subgroups (Figure 1). Our results also show that different regions of the flock tend to fly at different speeds; lead and edge birds are faster than middle birds (Table 3). Therefore, flocks are already shown to be dynamic in internal structure and may not have difficulty accommodating faster subgroups within the whole. Indeed, animals within groups frequently shift relative position during their movements (Cavagna, Queiros, Giardina, Stefanini, & Viale, 2013).

Prior research on pigeons found that flying in a cluster flock is energetically costly (Usherwood et al., 2011), either due to the additional maneuvering requirements to avoid neighbors or because of the turbulence produced by other birds. We also found evidence that, in addition to the potential aerodynamic benefit discussed above, shorebirds flying in a flock also suffer energetic costs. Specifically, birds flying further back in the flock and toward the edges flew more slowly and with increased wingbeat frequency. These effects are of the same order of magnitude as the aerodynamic neighbor term, suggesting that birds in aerodynamic position within the compound-V formation might be able to, on balance, escape the negative consequences of cluster flocks while birds out of position still experience them. If this is the case, it would indicate that the energetic cost of flying in a compound-V is between that observed in other cluster flocks and simple-V formations. However, other trends could also explain our results. For example, larger individuals are expected to have both lower flapping frequencies and faster flight speeds than smaller individuals of the same species. Thus, if larger individuals localize near the front of the flock, this could create the observed trends in airspeed and flapping frequency along the length of the flock.

In summary, our study reveals that four shorebird species of varying body size that have diverged evolutionarily for ~50M years employ a non-dimensional spacing rule that produces internal cluster flock relationships that mimic those of the simple-V formations of some larger migratory birds. Unlike cluster flocks in other species, these nearest-neighbor relationships also extend outward many meters and even affect whole-flock alignment, producing what we term a compound-V formation. Although the flight biomechanics data available in this study indicate a form of aerodynamic benefit in the compound-V flock, fully elucidating this as well as investigating the consequences for topics ranging from flock responsiveness to predators to migration remain topics for future research; progress in these areas is only likely to result from new theoretical modeling and data collection from on-bird loggers measuring physiological, flock positioning and biomechanical data. Nevertheless, our results indicate that a greater variety of bird flock structuring rules and internal relationships exist than has been revealed thus far. We propose that ecological demands are underappreciated drivers of diversity in the rules underlying animal collective behavior, and that further comparative studies will continue to reveal the diversity and importance of interaction rules in groups of animals.

## Materials and Methods

### Field Recording

We recorded multi-camera video of freely-behaving, wild birds in Humboldt county, California between April 17^th^-27^th^, 2017 and December 20^th^, 2017 to January 1^st^, 2018. Recordings were made at the Arcata Marsh Wildlife Sanctuary (40°51’25.35′N, 124° 5’39.37′W) and above agricultural fields in the Arcata bottoms (40°53’51.98′N, 124° 6’55.85′W). No birds were captured or handled, and we made efforts to avoid influencing bird behavior. Video was captured at 29.97 frames per second and 1920×1080 pixel resolution using three Canon 6D cameras with 35 mm or 50 mm lenses. Cameras were set along a 10 m transect and staggered in elevation. We setup cameras to overlook locations where birds aggregated during high tide or when foraging in agricultural fields. Flocking events included birds moving with the tide, or flushing in response to predators (e.g., peregrine falcons) or for unknown reasons. Cameras recorded continuously for up to 3 hours per day. For analysis, we selected flocks that included at least 100 individuals and that had an orientation and size allowing visual discrimination of individuals within the flock.

### Bird detection

We used the MATLAB R2017a (Natick, MA, USA) computer vision toolbox to generate code for detecting birds in video recordings. A foreground detector first separated moving objects from the stationary background. A gaussian filter was then applied to the image with a diameter matched to bird size under the recording conditions. Two-dimensional peak detection found local peaks in the smoothed image that were taken as potential bird positions.

Under some conditions, overlapping wings between adjacent birds prevented accurate detection of many individuals. To overcome this problem, we developed a frame-averaging algorithm that helped obscure the wings and emphasize the bodies. Here, optic flow determines the overall movement of the flock for each frame. Using the optic flow measurements and two-dimensional interpolation, the algorithm subtracts movement between frames. A rolling 5-frame window is then applied to the entire video. This procedure highlights pixels that are moving in the same direction as the flock, such as the birds’ bodies, while filtering pixels that are moving in other directions such as the wings.

### Three-dimensional calibration

Camera calibration followed established methodology (Hedrick, 2008; Jackson, Evangelista, Ray, & Hedrick, 2016; Theriault et al., 2014), with the exception that the distance between cameras was used to scale the scene instead of an object placed in view of the cameras. This approach allowed us to record in locations where it was infeasible to place calibration objects in front of the cameras (e.g., over water). The in-camera horizontal alignment feature was used to align cameras to the horizon. The pitch of the camera was measured with a digital inclinometer with 0.1-degree precision. This allowed alignment of the scene to gravity in post processing.

Background objects visible in the scene were used as calibration points. We developed a preliminary calibration using stationary objects such as trees, poles, and sitting birds. We then added flying birds, ensuring that points covered a wide range of distances and elevations relative to the cameras. Calibrations had low direct linear transformation (DLT) residuals (< 0.5-1 pixel), indicating high-quality calibrations.

### Camera synchronization

Cameras were synchronized by broadcasting audio tones over Walkie Talkies (Motorola Talkabout MH230) to each camera. Audio tones were broadcast approximately once every five minutes during recording. A time offset was determined for each pair of cameras using cross-correlation of the audio tracks. This offset allowed camera synchronization within ± one half of a frame, or 16.6 ms.

In recordings where birds were relatively close to the camera (< 50 m) and moving at relatively high pixel speeds, we used sub-frame interpolation to achieve increased synchronization accuracy of one tenth of a frame, or ± 1.7 ms. To determine the subframe offset, we interpolated tracks of moving birds used as background points in the calibration at 0.1 frame intervals from −1 to +1 frame (−1.0, −0.9, etc). We then calculated the DLT residual for a calibration with each combination of subframe-interpolated points for the three cameras. The set of offsets generating the lowest DLT residuals was used for the final calibration and applied to birds tracked in the study.

### Three-dimensional assignment

To reconstruct the three-dimensional positions of birds in a flock, 2D detections of individuals must be correctly assigned between cameras. We modified established software for this task (Evangelista et al., 2017; Zheng Wu, Hristov, Hedrick, Kunz, & Betke, 2009). Briefly, the software first finds all combinations of 2D points having DLT residual < 3 pixels. The software iteratively generates 3D points, starting with points having the lowest DLT residuals and only allowing a 2D detection to be reused a single time. This helps with the problem of occlusion while limiting the number of “ghost” birds (bird positions created from incorrectly matching detections among cameras). This process is repeated twice. The first iteration allows the user to determine a bounding region in 3D space where the flock is contained. In the second iteration, three-dimensional positions outside this bounding region are filtered before they can be considered as potential 3D points.

### Track generation

After 3D points have been generated, they are linked between frames to generate individual flight tracks. Here, a Kalman filter predicts the position of each bird in the subsequent frame for the 2D information from each camera and for the reconstructed 3D positions. In the first frame, the Kalman filter is seeded using optic flow measurements. For each frame step, a cost matrix is created from weighted sums of the 2D and 3D errors between predicted track positions and each reconstructed 3D point. The Hungarian algorithm is used to find a global optimum that minimizes the error in track assignment. A track that is not given an assignment is continued with a gap of up to 4 frames (0.13 s) after which it ends and any re-detection of the bird in question will start a new track.

### Wingbeat frequency analysis

We measured wingbeat frequencies in a subset of recordings where birds were both large enough and close enough to cameras to discern wingbeat oscillations. This excluded our smallest species, dunlin, and some flocks that were relatively distant from cameras. To measure wingbeat frequency, we used blob analysis to find a bounding box for each bird in each frame. We excluded blobs where the bounding box included two or more birds as determined using the track-assignment algorithm described above. We averaged four components of the bounding box to measure wingbeat phase: height, inverse of the width, detrended X-coordinate of top-left corner, and inverse of detrended Y-coordinate of the top-left corner. This allowed quantification of wingbeat phase independent of bird orientation with respect to the cameras. Wingbeat phase was averaged across cameras and bandpass filtered before a 128-point Fast Fourier Transform (FFT) was applied to measure wingbeat frequency. The frame rate of the cameras (29.97 frames per second) and the FFT window determined a wingbeat frequency bin size of 0.12 wingbeats s^−1^. Our method is similar to that used in a recent study of two corvid species (Ling et al., 2018).

### Species identification in mixed-species flocks

We recorded two mixed-species flocks of godwits and dowitchers. The size difference between species allowed species identification using the detected pixel area and distance of each bird (Figure 8). Here, blob analysis quantifies the pixel area for each bird in each tracked frame. Area data were excluded when two tracked birds were within a single blob bounding box. A low-pass filter was applied to the sequence of pixel area data across frames for each tracked bird to remove wingbeat effects. An object’s pixel area scales with the inverse of the square root of distance. Therefore, for each frame the square root of the filtered pixel area was multiplied by bird’s distance to provide a distance-scaled area. This value was averaged across frames and across cameras for each bird track. In mixed-species flocks a histogram of the scaled area had two distinct peaks with only a small amount of overlap (Figure 8a). Fitting two normal distributions to these data revealed an expected error rate in species identification of 3.3%. The scaled area where the two normal distributions intersect was used as the threshold for species identification.

**Figure 8.**
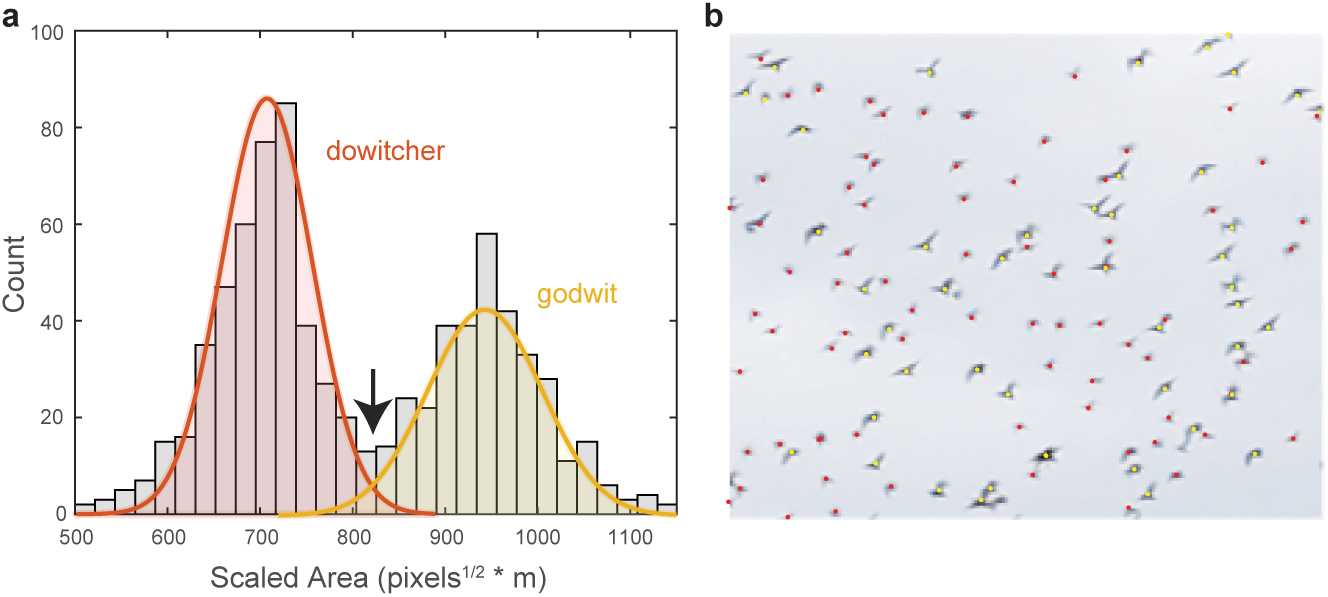
Species identification in mixed flocks. (a) Histogram of scaled pixel area of birds within a mixed-species flock. The two peaks are modeled as normal distributions. The area value where the two distributions intersect (indicated by the arrow) is used as the threshold for species identification. (b) Example section of a mixed-species flock with species identifications labeled by color.

### Neighbor alignment metrics

We quantified the relative position of each bird and its nearest neighbor in the flock (Figure 2). This was done separately for neighbors within ± 1 wingspan in flight elevation—the potential positions where aerodynamic interactions and collisions are plausible—and for neighbors beyond ± 1 wingspan. For each flock we calculated the modal lateral distance and modal front-back distance by taking the peak of a probability density function generated with a kernel density estimator and a smoothing bandwidth of 0.25 wingspans. We used modal values because distance calculations are truncated at zero, producing skewed distributions.

In a subsequent analysis (Figure 6), we quantified the angular distribution of neighbors at distances of 2, 4, 6, and 8 wingspans, and at a maximum radius depending on the size of the flock. Two-wingspan bins centered at the reference distance were used for selecting data points (e.g. birds within 1-3 wingspans were included in the 2-wingspan bin). Our aim was to examine the extent of internal structure within the flock. Because edge effects could create the appearance of internal structure, we excluded birds whose edge distance was less than the wingspan of the bin being analyzed. For example, for the 2-wingspan analysis, all birds within 3 wingspans of the horizontal edge of the flock were excluded. The maximum radius was taken as the median horizontal edge distance of all birds in the flock (Figure 9). This ensured that our analysis always included at least half of the flock.

**Figure 9.**
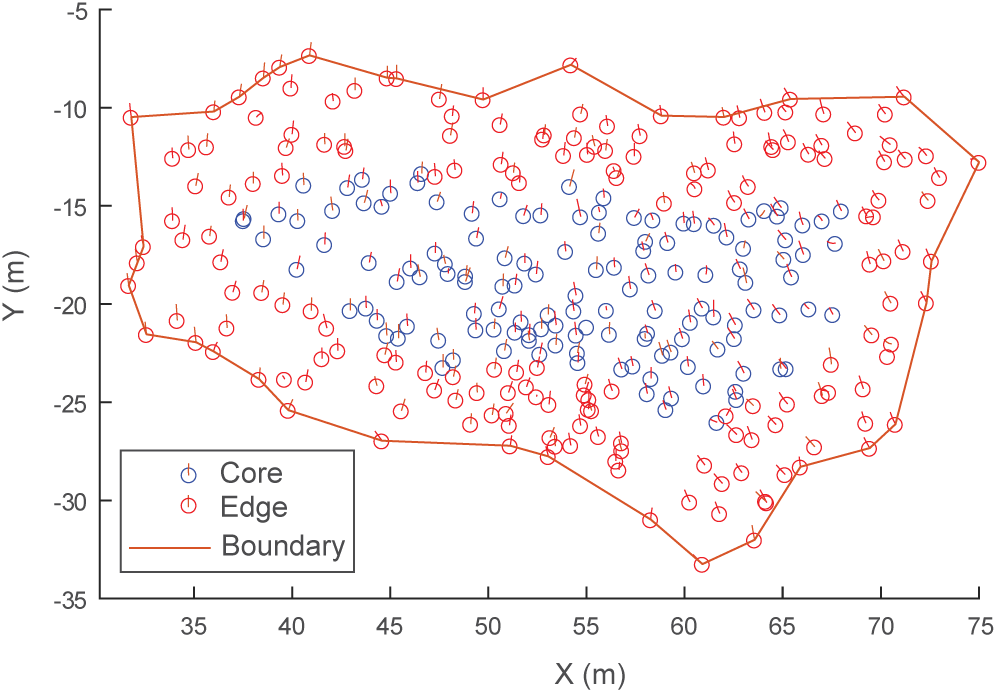
Determining flock edge and maximum radius. An overhead view of an example flock of avocets (flock 1220-2) is shown. Because flocks were always spread out in the horizontal direction, a compact hull is fit to the XY-coordinates to create a boundary. The minimum horizontal distance of each bird to the hull is the bird’s edge distance. The median edge distance is taken as the flock’s maximum radius for computing alignment metrics (Figure 6). Here, birds within the maximum edge distance (6.5 wingspans or 4.55 m) are labeled edge, and birds beyond the maximum edge distance are labeled core.

### Wingbeat phase analysis

We conducted an analysis to test for temporal and spatial wingbeat phase synchronization, following previously established methods (Portugal et al., 2014). We selected pairs of nearest neighbors in flocks where simultaneous wingbeat frequency data were available for both individuals for at least 20 frames (0.66 s). Cross correlation was used to determine the temporal phase offset between the birds. This value was divided by 2π*d*, where *d* is wingbeat duration, to attain a value between 0 and 2π. The spatial phase offset equals the temporal phase offset minus 2πλ, where λ is wingbeat wavelength. We tested for temporal and spatial synchrony by applying Rayleigh’s test for homogeneity of circular data to the temporal and spatial phase delays.

### Estimating wind speed and direction

We estimated local wind speed and direction for each flock using observed variation of ground speeds from birds flying in different directions. Ground speeds and flight directions were calculated for each bird at 1-wingbeat time intervals. Median ground speed was calculated for each 10-degree bin having at least 500 data points. A circle was then fit to these median values, with the center of the circle representing a vector of wind direction and magnitude. Ground speeds and wind direction and magnitude were then used to calculate airspeeds. This approach assumes that airspeed is independent of wind direction.

We compared our wind estimates to data from nearby weather stations. Our estimated wind direction was typically within ± 45 degrees and ± 2 m s^−1^ of weather station data (weather station KCAARCAT25). To avoid disturbing the birds, we did not attempt to release helium balloons to measure local wind conditions at altitude. Note that because our analysis here is based almost entirely on the positions and speeds of birds relative to their neighbors, our results are largely insensitive to the wind speed and direction.

### Statistical Analysis

Analyses were conducted using the statistical toolbox in MATLAB r2017b (The Mathworks, Natick, MA USA). We tested uniformity of circular distributions using Rao’s test(Fisher, 1995). Because multiple peaks were sometimes present, modal values were calculated using a circular kernel density estimator as an indicator of the predominant alignment direction. For the biomechanical analysis, we used linear mixed-effects models to predict individual wingbeat frequency and airspeed from 7 fixed effects—nearest neighbor distance, nearest-neighbor lateral distance, edge distance, airspeed, vertical speed, nearest neighbor species, front-back flock position and hypothesized aerodynamic positioning. Bayesian information criterion was used for model selection. All P-values were computed assuming two-tailed distributions.

## Data Availability

Source data for all figures and tables is included as part of this eLife submission.

## Acknowledgements

This work was funded by National Science Foundation grant IOS-1253276 to T.L.H. We thank Jonathan Rader for assisting with data collection and Jim Usherwood and William Conner for reviewing earlier drafts of the manuscript.

